# Massive genome decay and insertion sequence expansion drive the evolution of a novel host-restricted bacterial pathogen

**DOI:** 10.1101/2020.10.13.331058

**Authors:** Gonzalo Yebra, Andreas F Haag, Maan M Neamah, Bryan A Wee, Emily J Richardson, Pilar Horcajo, Sander Granneman, María Ángeles Tormo-Más, Ricardo de la Fuente, J Ross Fitzgerald, José R Penadés

**Author notes:** These authors contributed equally (Gonzalo Yebra, Andreas F Haag). Joint senior and corresponding authors. **Email addresses:** GY; AH. R de la F; JRF.

## Abstract

**Background:** The emergence of new pathogens is a major threat to public and veterinary health. Changes in bacterial habitat such as those associated with a switch in host or disease tropism are often accompanied by genetic adaptation. *Staphylococcus aureus* is a multi-host bacterial species comprising strains with distinct tropisms for human and livestock species. A microaerophilic subspecies, *Staphylococcus aureus* subsp. *anaerobius*, is responsible for outbreaks of Morel’s disease, a lymphadenitis in small ruminants. However, the evolutionary history of *S. aureus* subsp. *anaerobius* and its relatedness to *S. aureus* are unknown.

**Results:** Evolutionary genomic analyses of clinical *S. aureus* subsp. *anaerobius* isolates revealed a highly conserved clone that descended from a *S. aureus* progenitor about 1000 years ago before differentiating into distinct lineages representing African and European isolates. *S. aureus* subsp. *anaerobius* has undergone limited clonal expansion, with a restricted population size, and an evolutionary rate 10-fold slower than *S. aureus*. The transition to its current restricted ecological niche involved acquisition of a pathogenicity island encoding a ruminant host-specific effector of abscess formation, several large chromosomal re-arrangements, and the accumulation of at least 205 pseudogenes resulting in a highly fastidious metabolism. Importantly, expansion of ∼87 insertion sequences (IS) located largely in intergenic regions provided distinct mechanisms for the control of expression of flanking genes, representing a novel concept of the IS regulon.

**Conclusions:** Our findings reveal the remarkable evolutionary trajectory of a host-restricted bacterial pathogen that resulted from extensive remodelling of the *S. aureus* genome through an array of parallel mechanisms.

## Background

Bacteria have a remarkable capacity to adapt to new environmental niches, a characteristic that underpins their potential to become successful pathogens. Some pathogens can infect multiple host-species [1], while others become specialized and restricted to a single host. The evolutionary process of host restriction, observed across distantly-related bacterial groups, typically involves genomic events such as gene loss, gene acquisition, or chromosomal rearrangements [2-5]. The impact of such events may be most apparent after long-term associations between bacterium and host, with obligate intracellular pathogens the most extreme examples, exhibiting extremely small and compact genomes [6].

*Staphylococcus aureus* is a highly versatile bacterial pathogen, associated with an array of diseases in humans and livestock representing a threat to public and livestock health [7]. *S. aureus* has undergone extensive host-switching events during its evolutionary history leading to the emergence of new endemic and epidemic clones in livestock and humans [8]. In particular, the highest number of host-jump events appeared to have been between humans and ruminants in either direction [8]. A combination of gene acquisition, loss of gene function, and allelic diversification have been central to the capacity of *S. aureus* to undergo successful host-adaptation [8].

A close relative of *S. aureus, S. aureus* subsp. *anaerobius*, responsible for Morel’s disease [9], is highly restricted to small ruminants and its clinical manifestation is confined to abscesses located at major superficial lymph nodes. Outbreaks of this disease are associated with significant economic losses and have been reported in Europe, Africa and the Middle East [10]. In contrast to *S. aureus* subsp. *aureus, S. aureus* subsp. *anaerobius* is microaerophilic and catalase-negative [11], but their genetic and evolutionary relatedness is poorly understood.

Here, we investigate the evolutionary history of *S. aureus* subsp. anaerobius, revealing that it descended from an ancestor resembling *S. aureus* and adapted to its current ecological niche by extensive genome diversification involving massive genome decay, gene acquisition, and genome rearrangements. Furthermore, acquisition and expansion of ISSau8-like insertion sequences associated with the manipulation of transcription of flanking genes through multiple mechanisms leads us to propose the novel concept of the IS regulon. Overall, our results reveal a remarkable example of a bacterial pathogen that has undergone extensive genome remodelling in transition to a highly niche-restricted ecology.

## Results

### *S. aureus* subsp. *anaerobius* has a highly conserved genome

Previously, it has been reported that *S. aureus* subsp. *anaerobius* isolates belong to a single clonal complex as determined by MLST [10, 12]. To date only one fragmented draft genome of *S. aureus* subsp. *anaerobius* isolated in Sudan has been publicly available, and therefore the genome architecture has been unclear [13]. Here, we applied PacBio sequencing to the type strain MVF7, isolated in Spain in 1981-1982 [11], to provide the first complete genome sequence for a strain of *S. aureus* subsp. *anaerobius*. The genome consisted of a circular chromosome of 2,755,024 bp with 2,888 coding DNA sequences (CDS), 56 tRNAs and 5 complete rRNA copies, with a GC content of 32.74% (**Fig. 1A**).

**Figure 1:**
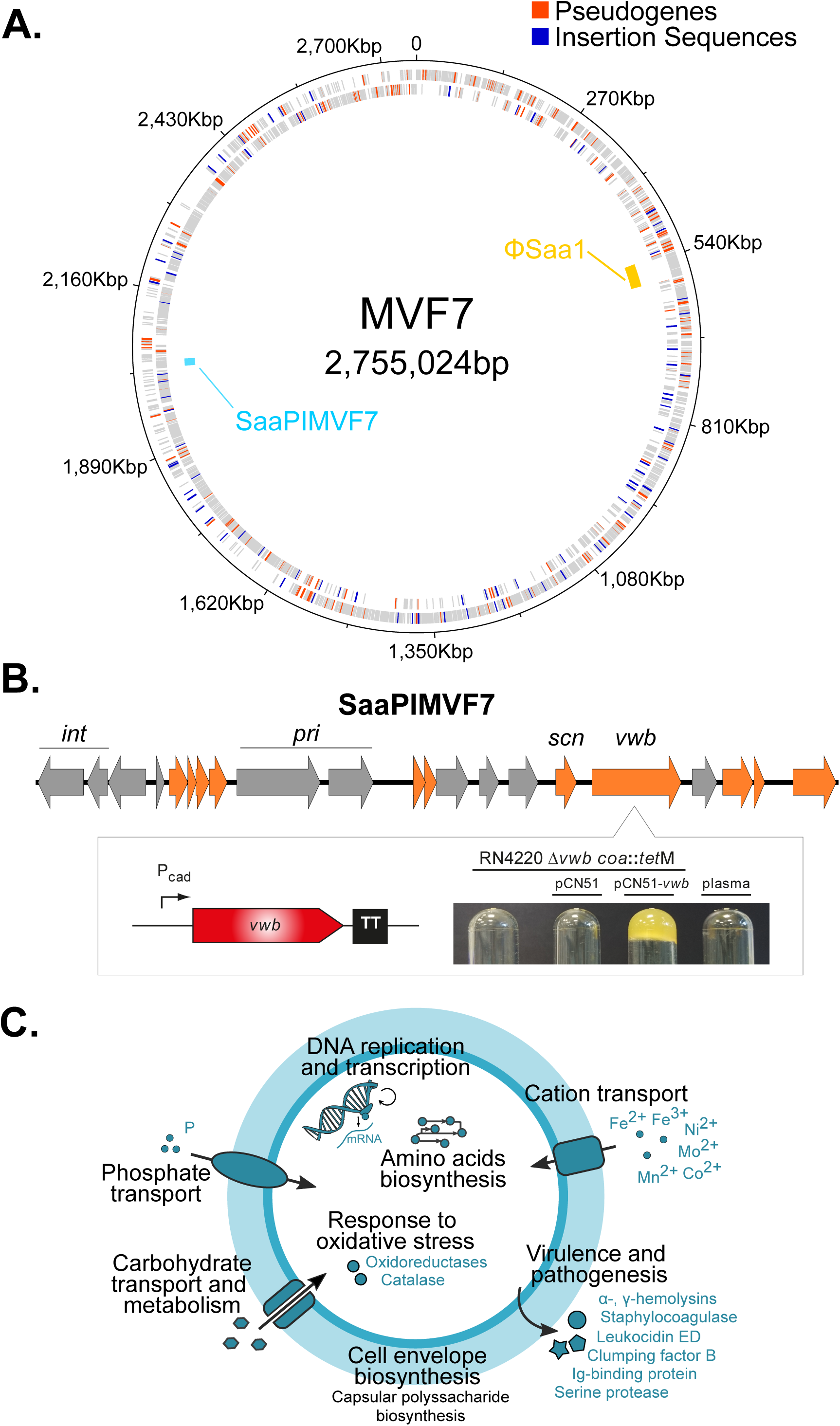
Pseudogenes, insertion sequences and mobile genetic elements in the *Staphylococcus aureus* subsp. *anaerobius* isolate MVF7. (A) Circular map of the chromosome. Rings show, from outside to inside: annotated genes in the positive strand, those in the negative strand (with the 205 pseudogenes and 87 insertion sequences shown in red and blue, respectively), and the mobile genetic elements found across all isolates (gold: prophage ΦSaa1; cyan: pathogenicity island SaaPIMVF7). (B) Gene map of SaaPIMVF7. Genes in grey are pseudogenes and genes in orange are intact by comparison other SaPI relatives. *int:* integrase; *pri:* primase; *scn*: Staphylococcal complement inhibitor (SCIN); *vwb*: von Willebrand factor-binding protein (vWBP). The box shows the expression of the SaaPIMVF7-encoded *vwb*. The SaaPIMVF7 *vwb* gene was cloned into the expression vector pCN51 under the control of a cadmium-inducible promoter, transformed into a coagulase and vWbp-deficient derivative of strain RN4220 (RN4220 *coa*::*tet*M Δ*vwb*) and the ability of SaaPIMVF7 vWbp to coagulate ruminant plasma was assessed. (C) Graphical summary of the main biological functions potentially disrupted by the presence of pseudogenes.

In order to further explore the genome content of *S. aureus* subsp. *anaerobius*, we obtained genomic DNA for 39 additional strains and performed Illumina whole genome sequencing to examine genome diversity and population genetic structure. Of these strains, 30 were isolated from different outbreaks in Spain between 1981 and 2012, with the others from Sudan (n=3), Italy (n=3), Poland (n=2) and Denmark (n=1) [10]. Among the isolates, we identified 4 closely-related novel STs (see **Additional Information** in Additional File 1 for more detail).

The combined dataset of 41 genomes had an average length of 2,685,366 bp (range: 2,604,446 to 2,758,945 bp) that contained 2,479 genes (92.7%) shared among all isolates (core-genome) of the total 2,675 genes identified (pan-genome). On average, each strain has a highly compact accessory genome of only 130 genes (range 107-143).

### *S. aureus* subsp. *anaerobius* evolved from an ancestor that resembled *S. aureus* subsp. *aureus* over a millennium ago

The evolutionary origin of *S. aureus* subsp. *anaerobius* and its phylogenetic relatedness to *S. aureus* subsp. *aureus* is unknown. In order to investigate its evolutionary history, we constructed a core genome alignment of *S. aureus* subsp. *anaerobius* and 807 *S. aureus* isolates that were representative of the species diversity [8] and reconstructed a maximum-likelihood phylogeny (**Fig. 2**). *S. aureus* subsp. *anaerobius* formed a distinct clade on a long branch within the *S. aureus* subsp. *aureus* diversity but was not genetically allied to any particular clonal complex. These data indicate that *S. aureus* subsp. *anaerobius* emerged from an unknown *S. aureus* lineage likely after a human to ruminant host-switch event that occurred over a millennium ago.

**Figure 2:**
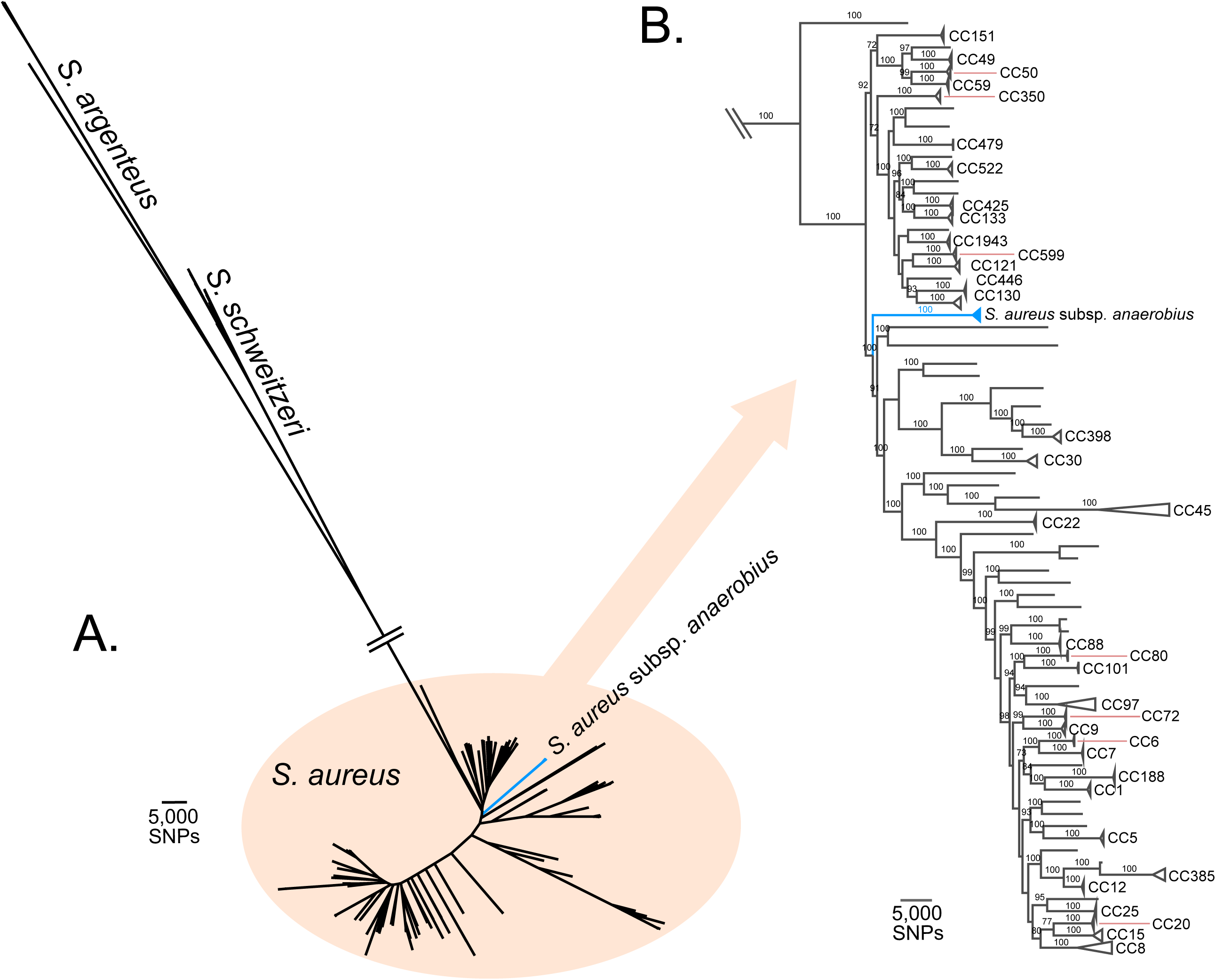
*Staphylococcus aureus* subsp. *anaerobius* represents a single clade within the *S. aureus* phylogenetic tree. Maximum Likelihood tree constructed from a SNP alignment of the studied *S. aureus* subsp. *anaerobius* sequences (clades in blue) and 787 sequences of different *S. aureus* subsp. *aureus* (in black). (A) Unrooted tree showing the divergence of 17 sequences of other *Staphylococcus* (*S. schweitzeri* and *S. argenteus*) whereas *S. aureus* subsp. *aureus* is embedded in the *S. aureus* diversity. (B) Subtree showing the position of the *S. aureus* subsp. *anaerobius* clade (collapsed in blue) with respect to the other *S. aureus* subsp. *aureus* clonal complexes (CC).

A core genome sequence alignment of the 41 isolates was produced with 5 recombinant regions (size range: 54 bp to 2,950 bp) excluded leaving an alignment of 2,324,010 sites with 3,443 variable sites. Phylogenetic analyses by the maximum likelihood approach revealed two distinct clades among the *S. aureus* subsp. *anaerobius* examined representing the isolates sampled in Sudan, and all others (sampled in Europe), respectively (**Fig. 3**). The average number of SNPs between pairs of *S. aureus* subsp. *anaerobius* genomes was 494 SNPs (range: 0-1,746), and the number of SNPs between the two clades was 1,697 SNPs (1,663-1,746).

**Figure 3:**
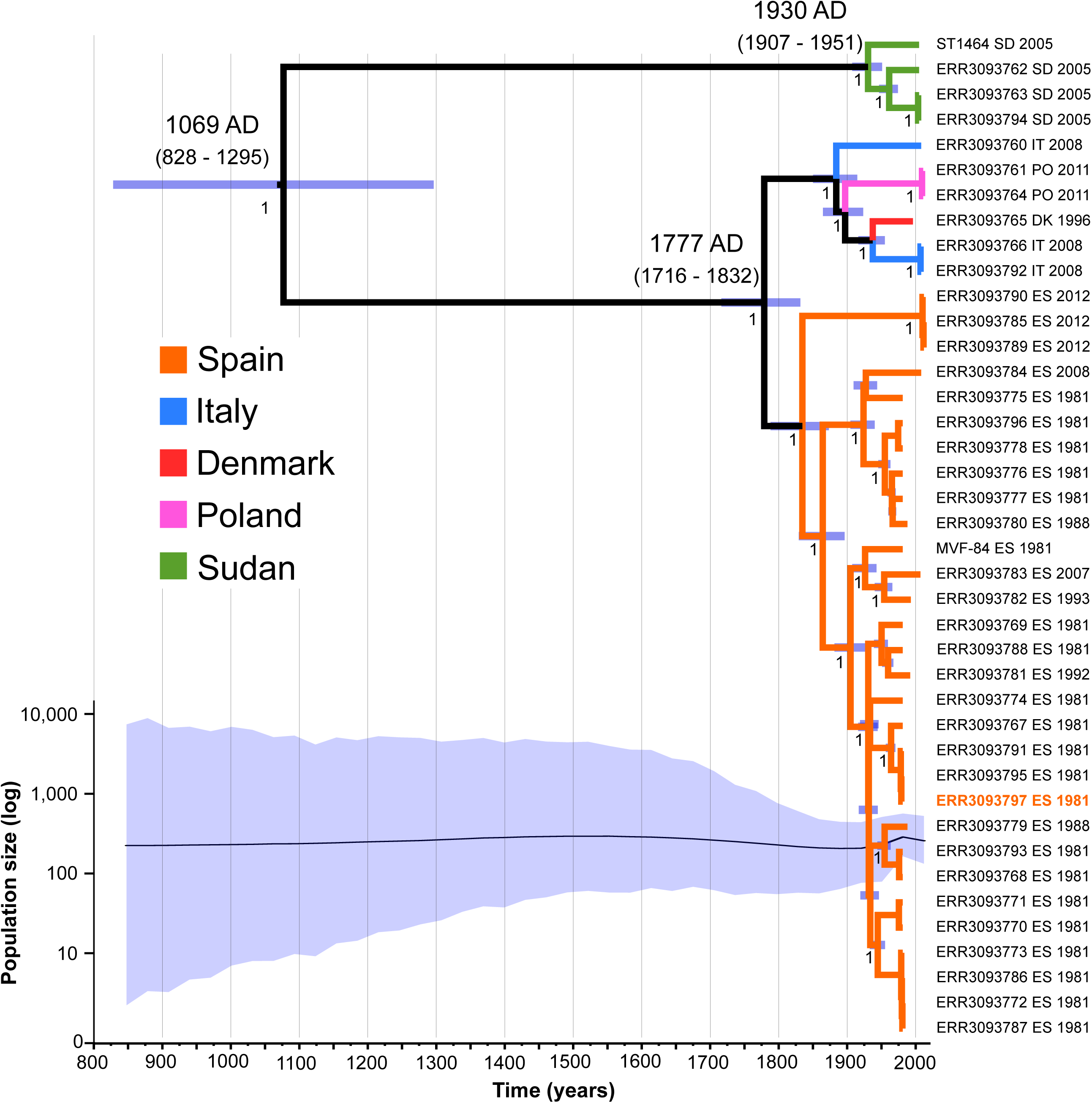
*S. aureus* subsp. *anaerobius* evolved over the last 1000 years with a stable population size. Bayesian maximum clade credibility tree of the *Staphylococcus aureus* subsp. *anaerobius* isolates. The x-axis is expressed in calendar years, branch colours denote sampling country. Purple bars at each node show its most recent common ancestor (MRCA) confidence interval. Numbers in nodes indicate the posterior probability. The estimated dates of the MRCAs of the main lineages are indicated. The isolate that was whole genome sequenced using PacBio technology (MVF7) is highlighted in orange. The bottom plot represents the changes in the effective population size over time, with the shadowed area representing the 95% credible interval, following the same timescale as the tree.

Linear regression analysis provided evidence of a molecular clock-like evolution in the ML phylogeny (**Fig. S1** in Additional File 1) and time-scaled Bayesian phylogenetic analysis indicated that the model combination that best fitted the data was a strict molecular clock paired with a constant population size, though other models performed similarly (data not shown). This analysis dated the most recent common ancestor (MRCA) of *S. aureus* subsp. *anaerobius* in 943 years before present (YBP) (95% Bayesian confidence interval (BCI): 1,184-716) (**Fig. 3**). The European and Sudanese clades diverged between 235 YBP (95% BCI: 296-180) and 82 YBP (95% BCI: 105-61), respectively. The estimated substitution rate was 4.4 (95% BCI: 3.3-5.5) × 10^−7^ substitutions/site/year (s/s/y), approximating to 1.2 SNPs per year which is ∼one order of magnitude slower than *S. aureus* subsp. *aureus* and may reflect the relatively slow growth observed for *S. aureus* subsp. *anaerobius* [9, 14].

### *S. aureus* subsp. *anaerobius* has undergone large intra-chromosomal rearrangements

During the evolutionary genomic analysis, we discovered that *S. aureus* subsp. *anaerobius* has undergone six large genomic translocations that ranged in size between 70 kb and 346 kb since separation from its *S. aureus* subsp. *aureus* progenitor (**Fig. 4** and **Fig. S2** in Additional File 1). As the edges of each translocated region were flanked by identical insertion sequence elements, we suggest that each event was the result of transposase-mediated homologous recombination between the leading and lagging strands [15]. For 2 pairs of translocations (I, VI, and II, V, respectively) the original genomic locations were exchanged whereas translocations III and IV were the result of excision and subsequent insertion at a distinct genomic location (**Fig. 4**). Because all translocations occurred via an inversion event, genes in the translocated areas conserved their original orientation and the GC skew remained unaffected (**Fig. 4**).

**Figure 4:**
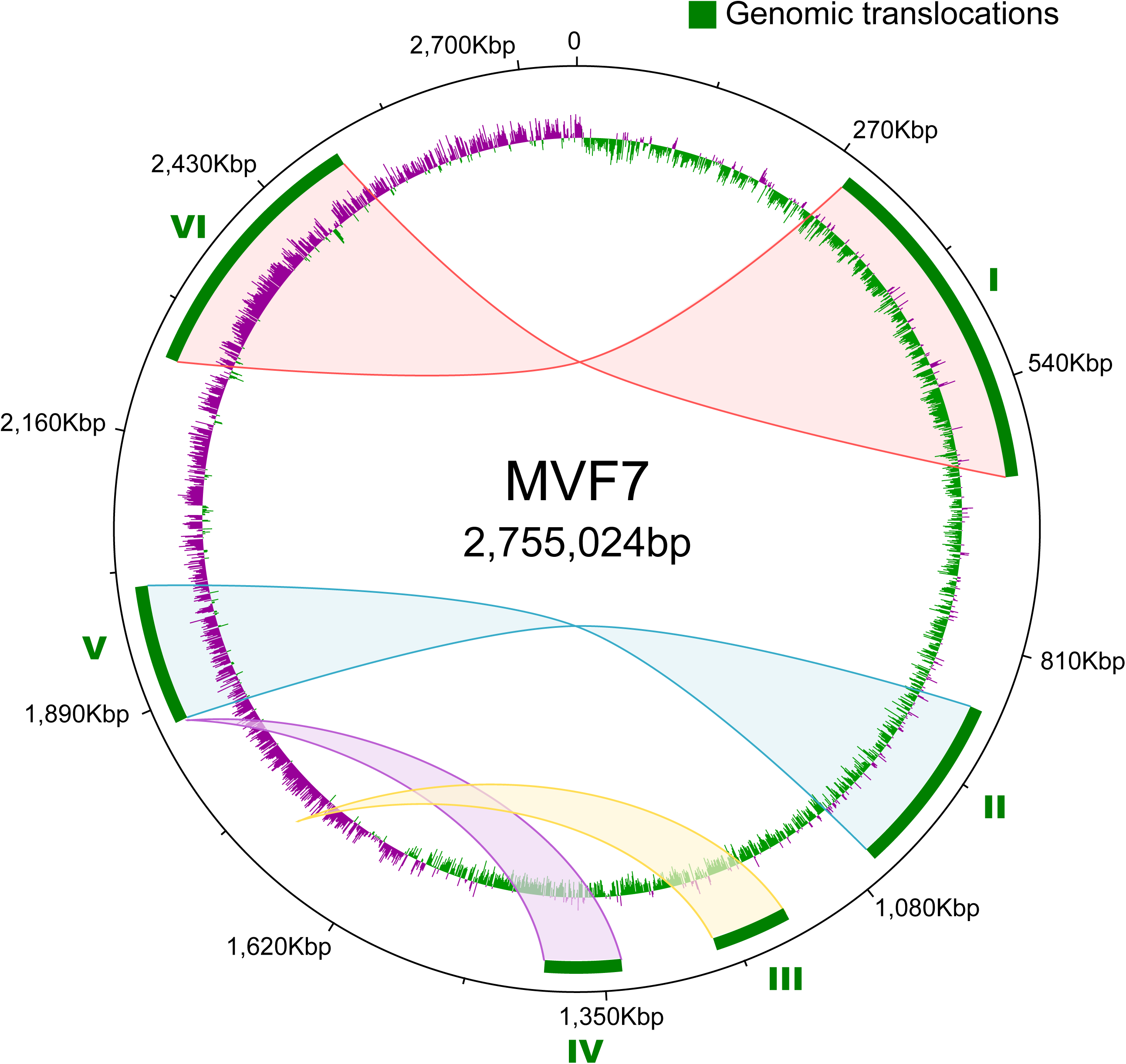
*S. aureus* subsp. *anaerobius* has undergone 6 large intra-chromosomal rearrangements. Map of the genomic translocations across the *Staphylococcus aureus* subsp. *anaerobius* isolate MVF7 chromosome. The green blocks represent the 6 large translocations detected in MVF7 when compared to the *Staphylococcus aureus* subsp. *aureus* RF122 strain (see also **Fig. S2** in the Additional File 1). The shaded areas indicate the original location of the translocated portions in the putative ancestral genome. The inner ring shows the GC skew (green for positive skew, purple for negative), which is unaffected by the translocations.

PCR-based analysis of the distribution of the 6 rearrangements identified in strain MVF7 among all *S. aureus* subsp. *anaerobius* strains revealed that all 6 were conserved in the majority of European isolates (clade II) whereas the isolates from Sudan (clade I) lacked translocations numbers II and V, suggesting they occurred in the clade I lineage since separation from a common ancestor (**Table S1** in Additional File 2). Although the impact of the re-arrangements on global gene expression is unclear, large chromosomal rearrangements in *S. aureus* subsp. *aureus* have been demonstrated to mediate transition to less virulent strains associated with persistent infection [16, 17].

### *S. aureus* subsp. *anaerobius* exhibits massive genome decay

Analysis of the complete genome of *S. aureus* subsp. *anaerobius* strain MVF7 revealed the existence of 205 pseudogenes, with a range of 201 to 210 pseudogenes per genome across the 41 *S. aureus* subsp. *anaerobius* strains examined (see **Additional Information** in Additional File 1). Pseudogenes originated through point mutations that caused frameshifts, premature stop codons or alternative downstream start codons and were evenly distributed around the genome (**Fig. 1A**). Relative to *S. aureus* subsp. *aureus*, (which contains approximately 20-50 pseudogenes per strain [18-20]), the high frequency of pseudogenes representing on average of 9.5% (range: 9.2-10.1%) of the total genome length represents a remarkable example of extensive gene loss that is likely to have a major impact on bacterial phenotype. A total of 164 pseudogenes (approximately 80% of those found in any given isolate) were shared among all *S. aureus* subsp. *anaerobius* genomes, of which 92.1% were caused by the same mutation in all isolates consistent with early events in the evolutionary history of *S. aureus* subsp. *anaerobius*.

The original functions of the 205 pseudogenes present in MVF7 were classified into clusters of orthologous groups (COGs) of proteins (see **Table S2** in Additional File 2). Of these, 82 (50%) were associated with metabolic functions, 29 (17.7%) to cellular processes and signalling, and 10 (6.1%) to information storage and processing, and 43 (26.2%) had unknown functions. Enrichment analyses of both KEGG pathways and GO terms (see **Table S3** in Additional File 2 for a full list) revealed a statistically significant enrichment of pseudogenes involved in biosynthesis of amino acids and metabolic pathways (see **Additional Information** in Additional File 1 and **Table S2** in Additional File 2). Of note, loss of function of genes mediating resistance to oxidative killing mechanisms such as catalase and other oxidoreductases may underpin the micro-aerophilic growth phenotype. In addition, an array of genes with known roles in pathogenicity including capsule biosynthesis, adherence and secreted proteins mediating interactions with the host, are predicted to be non-functional. Taken together, the extensive loss of gene function impacting on bacterial metabolism and pathogenicity is consistent with the highly fastidious growth requirements and defined disease tropism of *S. aureus* subsp. *anaerobius* (**Fig. 1C**).

### *S. aureus* subsp. *anaerobius* contains a novel pathogenicity island encoding a ruminant specific effector of abscess formation

Our genomic analysis revealed a small accessory genome with limited strain-dependent variation in gene content. However, 2 putative MGE were identified in all strains examined including a 43.2 kb, novel prophage (ΦSaa1) belonging to the *Siphoviridae* family, and a novel 13 kb *S. aureus* subsp. *anaerobius* pathogenicity island (SaaPIMVF7) (**Fig. 1B**). While ΦSaa1 contained no putative determinants of virulence, SaaPIMVF7 encodes novel variants of known virulence factors Staphylococcal complement inhibitor (SCIN) and the von Willebrand factor-binding protein (vWbp) (**Fig. 1B**) [18, 21]. A phylogenetic network of all members of the SaPI family constructed with SplitsTree v4.14.6 and the NeighborNet algorithm [22] (**Fig. S3** in Additional File 1) revealed that SaaPIMVF7 is genetically closest (93.1% nucleotide identity) to SaPIov2, found among isolates of *S. aureus* subsp. *aureus* CC133 which has a tropism for small ruminants (**Fig. 1B**). Of note, SaaPIMVF7 contained 10 pseudogenes out of a total of 21 genes identified, including the integrase and primase genes, suggesting that it can no longer be mobilized but is a stable feature of the chromosome. Of note, both the *vwb* and *scn* genes were intact and transcribed (data not shown), consistent with functionality (**Fig. 1C**). The vWBP has previously been demonstrated to promote coagulation of plasma, and is involved with abscess formation during invasive infection [23, 24]. In order to test the hypothesis that SaaPIMVF7-encoded vWBP was functional, we cloned the *vwb* variant into the expression vector pCN51, introduced this plasmid into a coagulase and vWbp-deficient derivative of strain RN4220 (RN4220 *coa*::*tet*M Δ*vwb*). Importantly, expression of the variant vWBP protein conferred the capacity for coagulation of ruminant plasma (**Fig. 1B**). In summary, all strains of *S. aureus* subsp. *anaerobius* contain a novel SaPI indicating an acquisition event that occurred over 1000 years ago prior to divergence of the 2 major subclades. Strikingly, the accumulation of loss-of-function mutations in genes required for mobilization indicate that SaaPIMVF7 is a stable element of the genome and functional expression of vWBP with ruminant coagulation activity suggests a potential role in abscess formation, a defining characteristic of Morel’s disease.

### Expansion and intergenic location of insertion sequences in the genome of *S. aureus* subsp. *anaerobius*

An additional striking feature of *S. aureus* subsp. *anaerobius* was the presence of at least 87 insertion sequences (IS) distributed around the genome in strain MVF7 (**Fig. 1A**). Each IS had 99.3% nucleotide identity with each other, and 97% identity with the previously described ISSau8 from bovine *S. aureus* strain RF122. The great majority (77 of 87) had premature stop codons that truncated the transposase gene whereas only 10 IS contained full-length intact transposase genes. The distribution of the IS elements identified in MVF7 among the rest of *S. aureus* subsp. *anaerobius* isolates was examined by mapping Illumina reads for each strain against the border between each IS and its flanking gene in the closed genome of strain MVF7 to identify reads overlapping this border. This analysis revealed that 68 out of the 87 MVF7 ISs (78.2%) were inserted at an identical location in all 41 *S. aureus* subsp. *anaerobius* isolates consistent with an ancient acquisition that occurred during the emergence of *S. aureus* subsp. *anaerobius*. An additional 7 IS elements were shared among all the European isolates, but were absent from isolates from Sudan (**Fig. 3**).

Previously, IS elements have been shown to contribute to loss of gene function or phase variation via direct insertion into CDS sequences [25]. However, all except one IS element were inserted in intergenic regions (the single exception was inserted into the *msb*A CDS), and we speculated that the non-random distribution of IS elements may influence the expression of the chromosomal genes located adjacent to the IS insertion site.

### IS affects *neighbouring* gene regulation in *S. aureus* subsp. *anaerobius* by disruption of promoter region and operon structure

Our *in silico* analysis of promoter and transcriptional terminator (TT) sequences indicated that each IS had a functional promoter but lacked a TT between the *tnp* gene and the downstream gene, supporting the idea that the *tnp* transcript could influence expression of the 3’-located gene. Furthermore, 2 putative TT were identified at different sites in the IS: TT1 is located in the 5’ region of the IS genes, and is bidirectional, while TT2 is located at the 3’ end region of the IS and is unidirectional in the antisense orientation (see **Fig. 5**). The putative role for the bidirectional TT1 would be to prevent the interference between transcripts initiated either from the IS or from the gene located upstream to the IS. By contrast, TT2 would prevent the impact that an antisense transcript originated from a gene located downstream of the IS could have on the expression of the IS transcript (**Fig. 5**). This organisation suggests a strategy that allows IS transcripts to interfere with the genes located 3’ to the IS, while blocking any interference that transcripts from flanking genes could have on the IS-derived regulatory transcript.

**Figure 5:**
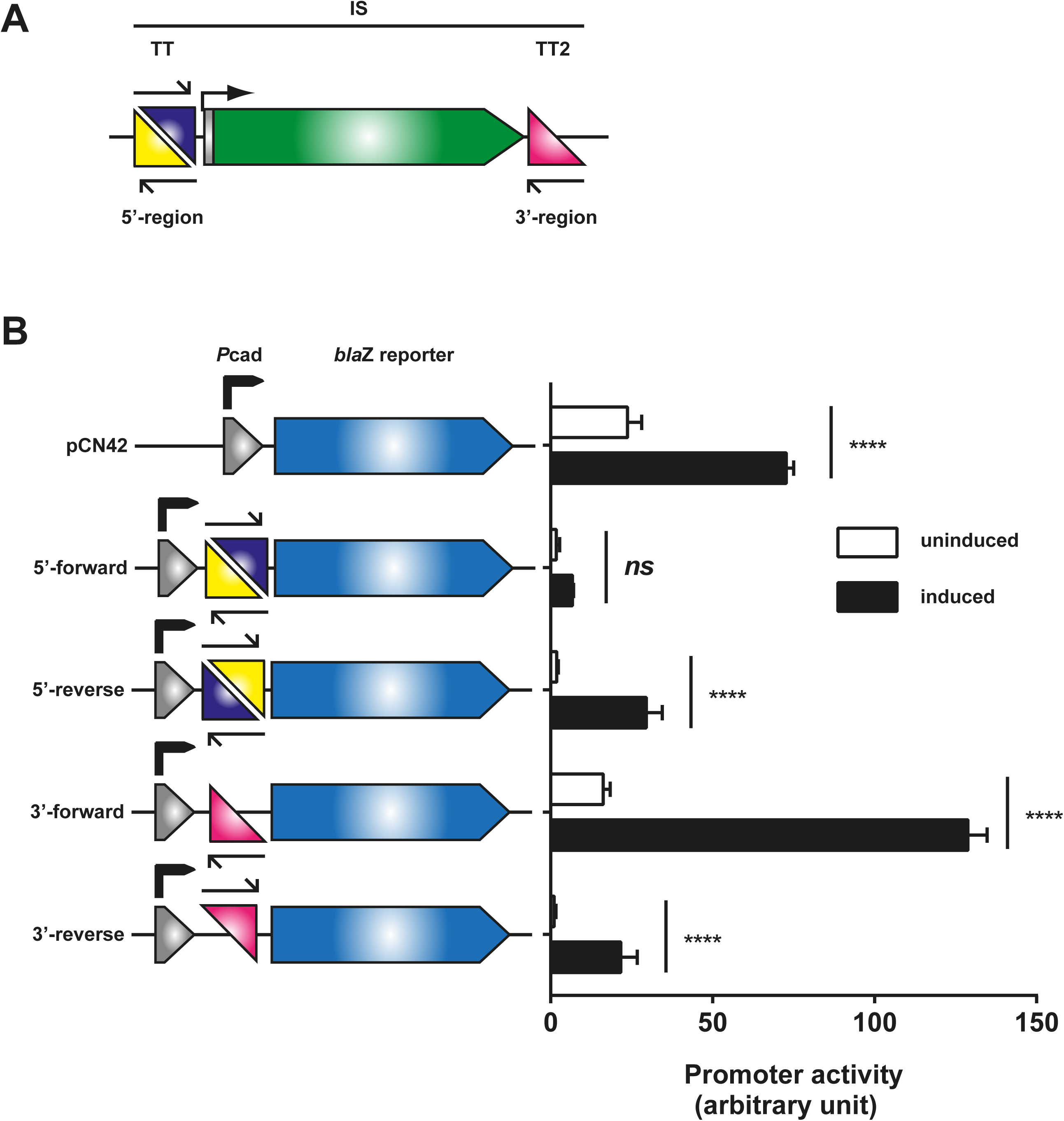
Characterisation of the transcriptional terminators (TTs) present in the ISs. (A) Schematic representation of IS-associated TTs. (B) Schematic representation of and functional assessment of the bi-directional TT1 localised in the 5’ region of the IS. The bi-directional TT1 is composed of two sub-TTs that share the same sequence but are inverted repeats, thus generating a master bi-directional TT1. The expression of the *bla*Z reporter in plasmid pCN42 is controlled by a cadmium-inducible promoter. The identified transcriptional terminators were cloned between the promoter and the *bla*Z reporter gene. β-lactamase expression was monitored either in uninduced (black bars) or Cd-induced (open bars) cultures 180 min after induction. Data are show the means of three independent biological replicates and error bars show the standard deviation from the mean. Statistical analysis was performed using two-way ANOVA followed by Dunnett’s post-test. Multiplicity adjusted p-values are shown for uninduced to induced comparison and all terminator containing reporters showed significant differences when compared to their respective empty plasmid control. **** p<0.0001, ns not significant.

To validate the functionality and directionality of the TTs, we employed plasmid pCN42 [26], in which the *bla*Z reporter is under the control of the P_*cad*_ promoter. The conserved 5’ and 3’ region of the ISs containing the bi- and unidirectional TTs, respectively, were inserted, in both orientations, between the P_*cad*_ promoter and the *bla*Z reporter (see **Fig. 5**), and the expression of the reporter gene measured with and without induction of the P*cad* promotor. The results confirm the existence, functionality and directionality of both TTs, consistent with the hypothesis that ISs can control the expression of 3’-located chromosomal genes.

To test the idea that the intergenic IS insertions arise by selection because they influence the expression of the neighbouring genes, we examined the impact of IS on the expression of selected downstream genes located in the same orientation as the IS. Since *S. aureus* subsp. *anaerobius* is not genetically tractable, we designed reporter constructs reconstructing the genetic organisation found in *S. aureus* subsp. *anaerobius* (**Fig. S4** in Additional File 1) and tested gene expression levels in *S. aureus* subsp. *aureus*. Specifically, we selected four different genes (*rps*P, *dna*D, *met*C and *met*N) that were identified in *S. aureus* subsp. *anaerobius* to contain an IS at distinct distances upstream of the gene start codon, and constructed *bla*Z transcriptional reporter fusions using pCN41 [26] (**Fig. 6A-D**). The expression of these genes in the presence or absence of upstream ISs was then tested. Importantly, and in support of the hypothesis, absence of an IS impacted on the expression of the neighbouring genes, confirming their regulatory role, although this effect was different depending on the gene including increased transcription of *dna*D, *met*C or *met*N, or decreased *rps*P transcription (**Fig. 6A-D**). The impact on downstream gene expression likely depends on the position of the IS relative to the gene’s start codon and the nature of its integration (conservative or deleterious; see **Fig. S4** in Additional File 1). In some cases, the IS will reduce expression by altering either the promoter or the DNA binding domain of a putative inducer of the flanking gene, while in other cases it may promote enhanced transcription by eliminating the binding site of a repressor. These hypotheses are currently under investigation.

**Figure 6:**
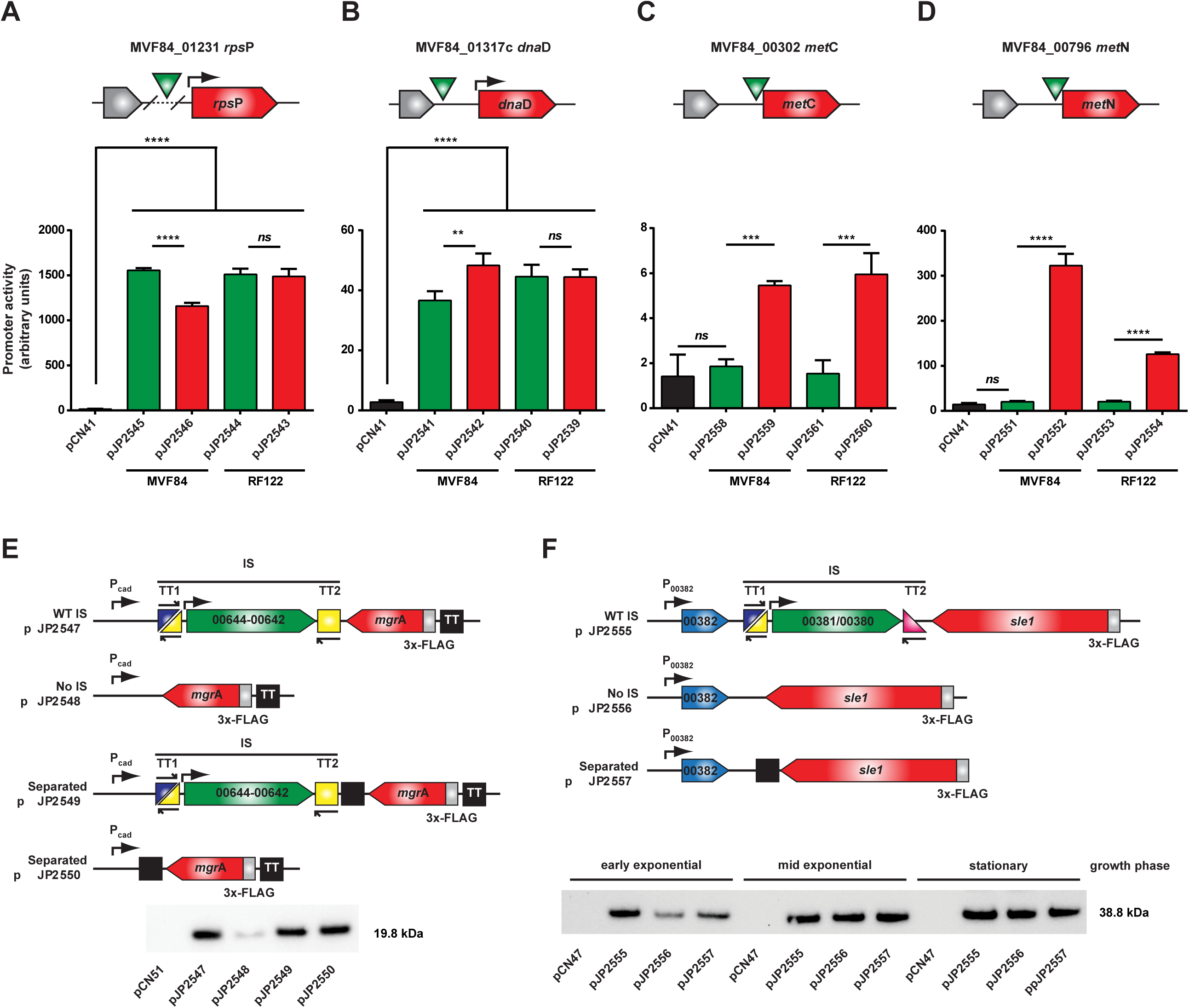
Presence of IS influences the expression of downstream genes and genes product through multiple mechanisms. (A-D) Reporter constructs for presence of the IS upstream and in sense orientation of the target gene were designed in plasmid pCN41 containing a β-lactamase reporter gene. The green triangle indicates the IS position relative to the target gene’s start. All reporters were constructed either by removing the IS gene from *S. aureus* subsp. *anaerobius* strain MVF84 or by introducing the IS into *S. aureus* subsp. *aureus* strain RF122 to confirm the IS role on gene expression variation. In (A) the IS removal results in a 207bp deletion to the promoter region not present in RF122. Reporter plasmids were introduced into RN4220 as described in Methods. Data show the mean of three biological replicates, error bars represent the mean’s standard deviation. One-Way ANOVA was performed followed by Tukey’s multiple comparisons test. *p<0.05, **p<0.01, ***p<0.001, ****p<0.0001, *ns* not significant. (E&F) Western blot analysis of (E) *mgr*A and (F) *sle1* expression constructs to address whether the IS impacts on their expression levels by interrupting expression of an antisense transcript. (E) *mgr*A encoding a 3xFLAG tag was cloned in antisense to a cadmium-inducible promoter into plasmid pCN51 to mimic the expression of an antisense transcript to *mgr*A. Constructs presented different combinations of presence/absence of IS and/or transcriptional terminator the IS transcript or expression from the plasmid-encoded cadmium-inducible promoter from the *mgr*A transcript. Expression of the cadmium-inducible promoter was induced with 5 µM CdCl_2_. (F) To assess the IS impact in the context of a natural antisense transcript, 3xFLAG-encoding expression constructs of *sle1* including the gene downstream of *sle1* in the ancestral genome were cloned into the plasmid pCN47. Samples were taken during different growth phases. Plasmids were introduced into RN4220 Δ*spa* as described in Methods. Western blots shown are representative of at least two independent biological replicates. TT, transcriptional terminator.

### ISs affect *S. aureus* subsp. *anaerobius* gene expression by a novel anti-antisense RNA interference mechanism

Intriguingly, the majority of ISs (63 of 87) were located in an antisense orientation to putative target genes. In such an orientation, the previously described effects on gene expression would not be feasible, and we reasoned that distinct mechanisms of IS-mediated control of neighbouring gene expression may exist. To test this hypothesis, we selected 2 examples of *S. aureus* subsp. *anaerobius* genes that were located in an anti-sense orientation to the flanking IS, including *mgr*A, encoding a global regulator of virulence [27-29], and *adh*A encoding a zinc-dependent alcohol dehydrogenase, induced under low oxygen conditions [30] (**Figs. S4** and **S5** in Additional File 1). Since the antisense orientation of these genes relative to IS precludes monitoring of gene expression by transcriptional fusions, they were cloned to express 3xFLAG tagged versions of the encoded proteins, which facilitated measurement of expression levels by western immunoblot. To examine the impact of IS on gene expression, expression plasmid constructs with and without IS in the antisense orientation were introduced to *S. aureus* subsp. *aureus* strain RN4220 Δ*spa* and expression levels compared (see scheme in **Fig. S5** in Additional File 1). Surprisingly, no differences in the expression levels of MgrA or AdhA were observed in any of the constructs suggesting that the presence of ISs did not actively alter expression levels of the downstream genes by a classical anti-sense mechanism (**Figs. S5A** and **S5B**, respectively, in Additional File 1).

However, the analysis of the genetic context of *mgr*A revealed another possible mechanism of IS-mediated control of gene expression. In the ancestral *S. aureus* subsp. *aureus*, the gene downstream of *mgr*A is in an antisense orientation (**Fig. S4** in Additional File 1). Furthermore, overlapping antisense transcripts for *mgr*A and its downstream gene in *S. aureus* have been identified in a previous study [31], suggesting the possibility of antisense interference of *mgrA* gene expression. Accordingly, we hypothesized that the presence of an IS between the 2 genes in *S. aureus* subsp. *anaerobius* could disrupt the putative interference (antisense) mechanism. To examine this possibility, we constructed another set of reporters (see scheme in **Fig. 4E**) in which we used the inducible expression plasmid pCN51 to mimic expression from the gene adjacent to *mgr*A. Since the gene located downstream of *mgr*A belongs to a large operon that was impossible to clone into pCN51, we used the cadmium-inducible promoter present in plasmid pCN51 as the origin of the anti-sense mRNA for *mgr*A, mimicking the natural genetic context (**Fig. 4E**). The different constructs were introduced into the *S. aureus* subsp. *aureus* strain RN4220 Δ*spa* and expression from the cadmium-inducible pCN51 promoter induced. Absence of the IS resulted in much reduced MgrA expression levels suggesting that presence of a strong transcriptional terminator protects *mgr*A mRNA from antisense mRNA interference (**Fig. 4E**). In support of this idea, insertion of a TT between *mgr*A and the plasmid-encoded cadmium-inducible promoter produced the same effect as the presence of the complete IS (**Fig. 4E**). A similar effect was identified on *sle1* encoding an N-acetylmuramyl-L-alanine amidase involved in staphylococcal cell separation and β-lactam resistance [32] [33], whereby IS insertion has disrupted the anti-sense regulation of Sle1 expression by a neighbouring gene in a growth phase-dependent manner (**Fig. 4F**). Thus, IS in *S. aureus* subsp. *anaerobius* can uncouple genes from ancestral antisense regulation by acting as anti-antisense elements, a hitherto unknown regulatory role.

Overall, our data suggest that the non-random insertion of IS into *S. aureus* subsp. *anaerobius* intergenic regions has influenced the expression of neighbouring genes through multiple distinct mechanisms. The putative attenuated expression of up to 87 different chromosomal genes involved in an array of different functions (**Table S4** in Additional File 2) supports the novel concept of the *IS regulon* in which the expression of a subset of genes in a bacterial genome is controlled, not by a transcriptional regulator, but by an IS.

## Discussion

While S. *aureus* is an opportunistic pathogen responsible for an array of different pathologies in different anatomical sites in humans and animals, *S. aureus* subsp. *anaerobius* is restricted to a specific infection of superficial lymph nodes in sheep and goats. Here, we provide a remarkable example of the evolutionary transition of a versatile multi-host bacterium to a fastidious highly niche-restricted endemic pathogen of small ruminants. The transition was marked by multiple distinct evolutionary processes mediating drastic changes to the genome that resulted in a complete re-modelling of bacterium-niche interactions. Previous studies have indicated a human ancestral host for *S. aureus* and endemic livestock clones are the result of host jump events that have occurred during the evolutionary history of *S. aureus* [8, 34]. Accordingly, the co-segregation of all isolates of the *S. aureus* subsp. *anaerobius* within a single monophyletic clade in the *S. aureus* phylogeny suggests a likely human to ruminant host-switch event that preceded the evolutionary transition to a highly niche-specific ecology.

Such switches to a host-restricted lifestyle [35], have occurred across the bacterial kingdom in diverse lineages such as *Mycobacterium* [36], *Shigella* [37], *Salmonella* [38], and *Burkholderia* [39], and extreme examples are represented by endosymbionts that have evolved from free-living organisms to become dependent on a single host species [2, 6]. While most known examples of host-restrictive evolution have time-scales of hundreds of thousands up to many millions of years [6, 35], our analysis indicates that *S. aureus* subsp. *anaerobius* evolved about 1000 or more years ago offering a unique insight into the relatively early stages of evolution towards niche-restriction.

Host shifts are associated with a radical change in habitat, typically with a genetic bottleneck that diminishes effective population size and a corresponding reduction in purifying selection activity [2, 35]. We discovered that approximately 10% of the *S. aureus* subsp. *anaerobius* genome is made up of pseudogenes affecting an array of metabolic and pathogenic pathways associated with the fastidious nutritional requirements and limited virulence of *S. aureus* subsp. *anaerobius* (**Fig. 1C**). In particular, the microaerophilic metabolism, is likely to be due to the loss of function of catalase and other oxidoreductase genes leading to increased sensitivity to oxidative free radicals. These data suggest adaptation to a nutrient-rich, oxygen-limited niche, such as that provided by the lymphatic system [11]. The repair of mildly deleterious mutations such as those resulting in loss of gene function is likely compromised by lower levels of purifying selection leading to fixation by genetic drift, and sufficient time has not elapsed to facilitate the deletion of genes no longer required in the new habitat, possibly also impacted by the observed lower rate of mutation.

We identified a large number of closely-related IS elements distributed around the genome of *S. aureus* subsp. *anaerobius* (**Fig. 1A**; **Table S2** in Additional File 2), a phenomenon that has been observed among other bacteria evolving towards host-restriction [2, 4, 6, 39]. Ineffective purifying selection after a population bottleneck also allows insertion sequences (IS), normally present in bacterial genomes in low numbers (<10), to expand and disseminate around the genome in the early stages of host-restriction. Over time the IS elements will be eventually purged, such that recently host-restricted bacteria will generally contain more IS than ancient symbionts [2, 4]. It is well established that IS elements often insert into coding regions resulting in gene inactivation and can also mediate chromosomal rearrangements of genetic segments flanked by ISs via homologous recombination events [15]. We identified five large rearrangements that occurred in the common ancestor of all isolates and an additional event exclusive to the clade comprising European isolates. Although the effects on bacterial phenotype resulting from the identified rearrangements is unknown, large rearrangements can change the distance of a gene from the origin of chromosome replication leading to altered gene copy number and expression [40, 41] and similar rearrangements in *S. aureus* have been reported to mediate transition to reduced virulence phenotypes such as single colony variants associated with persistent infections [16, 17].

While IS elements often inactivate genes, it is striking that only one of 87 IS insertions in *S. aureus* subsp. *anaerobius* results in gene disruption. It is established that IS elements can also activate expression of neighbouring genes, either by an extended transcription from an internal promoter or by the generation of a hybrid promoter [25, 42]. Uniquely, the IS-mediated gene regulation of *S. aureus* subsp. *anaerobius* involves one established and one novel mechanism of control. Upstream IS elements oriented in the sense orientation relative to the flanking gene (∼25% of IS) act mainly through modulation of promoter and operon structure as previously reported [43]. However, in *S. aureus* subsp. *anaerobius*, the majority of ISs (∼75%) are located downstream in the antisense orientation. In the ancestral *S. aureus* subsp. *aureus* strain, the expression of some genes is controlled by transcripts of a downstream gene in the antisense orientation suggesting expression of the corresponding proteins is mutually exclusive [44]. Here, we have shown that in *S. aureus* subsp. *anaerobius*, IS inserted in an antisense orientation to flanking genes can effectively uncouple the targeted genes from their interdependent expression control and act as anti-antisense regulatory elements. This novel regulatory mechanism might provide a selective advantage to *S. aureus* subsp. *anaerobius* in facilitating the simultaneous expression of proteins required concurrently the new niche.

While some bacterial pathogens such as *Bordetella pertussis* utilise IS elements to control the expression of their flanking genes in a strain-specific manner [43], the genomic localisation of the majority of IS elements in *S. aureus* subsp. *anaerobius* is conserved across the phylogeny, indicating that insertion events happened early in the evolution of *S. aureus* subsp. *anaerobius*, and suggesting that the complement of genes affected may be important for the ecology of *S. aureus* subsp. *anaerobius*. Based on this, we propose the novel concept of the *IS regulon*, representing the set of genes that is transcriptionally controlled by IS elements.

In conclusion, using a combination of phylodynamic, comparative genomic and molecular biology approaches, we have dissected the relatively recent evolution of a host-restricted bacterial pathogen underpinned by drastic changes to the genome via numerous distinct processes. In particular, IS elements have been domesticated by *S. aureus*, contributing to the chromosomal architecture via intra-chromosomal rearrangements, and control of gene expression via multiple mechanisms. Taken together, our findings provide a unique and remarkable example of the capacity of bacterial pathogens to expand into new host niches.

## Methods

### Bacterial strains and growth conditions

The bacterial strains employed are detailed in **Table S5** in Additional File 2. *S. aureus* was grown in Tryptic soy broth (TSB) or on Tryptic soy agar and *Escherichia coli* was grown in Luria-Bertani broth (LB) or on LB agar. Antibiotic selection was used where appropriate (erythromycin 10 µg ml^-1^ for *S. aureus* and ampicillin 100 µg ml^-1^ for *E. coli*.

### Whole genome sequencing and genomic analysis

Forty *S. aureus* subsp. *anaerobius* ovine isolates previously reported [10] were selected, of which 31 were sampled in Spain during different outbreaks spanning 3 decades (1981-2012). The rest of the isolates, sampled between 1996 and 2011, were from Sudan (n=3), Italy (n=3), Poland (n=2) and Denmark (n=1). Isolates were sequenced using a MiSeq machine (llumina, San Diego, CA, USA); the paired-end short reads were trimmed using Trimmomatic v0.36 [45] and *de novo* assembled into contigs using SPAdes v3.10.0 [46] and Velvet v1.2.10 [47]. Illumina read data is available in the European Nucleotide Archive under the study accession number PRJEB30965.

One of the Spanish isolates (MVF7, the type strain) was sequenced on the RSII platform (Pacific Biosciences, Menlo Park, CA, USA) using SMRT technology in the Centre for Genomic Research (University of Liverpool, UK). The long reads were de novo assembled into a single contig using the Hierarchical Genome Assembly Process (HGAP) method. Also included in the analysis was the only *S. aureus* subsp. *anaerobius* draft genome publicly available, isolate ST1464 (assembly GCA_000588835.1) [13]. Genes of all 41 genomes and draft assemblies were annotated using Prokka v1.12 [48], and the pan-genome was determined using Roary v3.8.2 [49] applying a 95% identity cut-off. *In silico* multilocus sequence typing (MLST) of the assemblies was performed using the MLST tool v2.8 (https://github.com/tseemann/mlst). Phage sequences and genomic islands were identified using PHASTER [50] and Island Viewer v4 [51], respectively, and manually inspected for reliability. Antimicrobial resistance genes were detected using ResFinder v3.0 [52].

### Pseudogene detection

Here, pseudogenes were defined as protein-coding sequences with homologous genes in an *S. aureus* subsp. *aureus* reference that were split or truncated (i.e. <80% of the reference gene length). For this purpose, the isolate RF122 (assembly GCA_000009005.1) was employed as a reference and a custom python script (https://github.com/GonzaloYebra/anaerobius) was developed to identify pseudogenes from the output of a Roary analysis of *S. aureus* subsp. *anaerobius* isolates compared to the *S. aureus* RF122 genome. The analyses were repeated using multiple *S. aureus* subsp. *aureus* genome sequence references, but this did not alter significantly the pseudogenes detected (data not shown). Ancestral functions of pseudogenes were predicted by assigning them to clusters of orthologous groups (COGs) using eggNOG v4.5.1 [53]. Enrichment analyses of GO (Gene Ontology) terms and KEGG (Kyoto Encyclopedia of Genes and Genomes) pathways assigned by eggNOG were performed using the R packages topGO v2.34.0 and ClusterProfiler v3.10.0 [54]. Additionally, the Integrated Microbial Genomes (IGM) annotation tool [55] was used to infer phenotypic metabolic characteristics from the presence/absence of protein pathways.

### Insertion sequence detection

Transposase coding sequences were identified in the MVF7 complete genome using the Prokka output and the ISsaga web tool hosted in the ISfinder platform [56]. Determining the location of IS elements and other repetitive sequences by assembly of short-read sequences is often not feasible [57]. Therefore, in order to examine the distribution of IS identified in strain MVF7 among other isolates, Illumina reads were mapped to fragments of the MVF7 genome representing each transposase edge together with their flanking genes using BWA-MEM [58]. In this manner, identification of reads that spanned the border between the IS and the flanking gene indicated the presence of that IS at the same genomic location relative to MVF7.

### Phylogenetic analyses

A core genome alignment was created by aligning the Illumina short reads and the ST1464 assembly to the MVF7 whole genome using Snippy v3.1 (https://github.com/tseemann/snippy). Sites containing any gap character (‘-’) or unknown nucleotide (‘N’) were discarded. Gubbins v2.2.0 [59] was used to detect recombinant regions which were then discarded. A maximum likelihood (ML) tree was constructed using IQ-TREE [60], applying the GTR nucleotide substitution model together with a gamma-distributed rate heterogeneity across sites and 1,000 ultrafast bootstrap replicates. Temporal signal was investigated using TempEst v1.5 [61] by means of the correlation between root-to-tip distances and sampling dates (**Fig. S1** in Additional File 1). Bayesian phylogenetic analysis was performed with BEAST v1.9.0 [62] using the HKY model for nucleotide substitution. Different models were tested for the molecular clock (strict and uncorrelated lognormal relaxed) and demographic (constant, exponential and Bayesian skygrid) models. Each of these model combinations were run for 100 million generations, with sampling every 10,000 and discarding the initial 10% as burn-in. Runs were compared via a marginal likelihood estimation (MLE) using path sampling and stepping stone sampling methods implemented in BEAST. The posterior distribution of trees was summarised into a maximum clade credibility tree.

In order to examine the relatedness of *S. aureus* subsp. *anaerobius* and *S. aureus*, another core genome SNP tree was created, also using Snippy v3.1 and IQ-TREE (same settings than above). It included a sample of 790 *S. aureus* genomes (corresponding to 43 different host species and 77 clonal complexes (CCs), isolated in 50 different countries) along with 17 isolates of the most closely-related staphylococcal species (*S. schweitzeri* and *S. argenteus*) [8]. All bioinformatic analyses were carried out using the Cloud Infrastructure for Microbial Bioinformatics (CLIMB) facility [63].

### Gene cloning

General DNA manipulations were performed using standard procedures. PCR fragments were amplified from genomic DNA using Kapa Hifi DNA polymerase (Kapa Biosystems). PCR fragments were either digested using restriction endonucleases and ligated into the respective plasmid backbone or assembled directly into a linearised plasmid backbone using Gibson assembly (NEBuilder HiFi DNA Assembly, NEB) according to the manufacturer’s instructions. The plasmids and oligonucleotides used in this study are listed in **Tables S6** and **S7** in Additional File 2, respectively.

### Western blot analysis

*S. aureus* cultures harbouring the respective pCN47 or pCN51 derivative plasmids were diluted 1:50 from overnight cultures and grown in TSB at 37°C and 120 rpm until sample collection. For pCN51 constructs induced with CdCl_2_, cultures were split at early exponential phase (OD_540_∼0.15), 5 μM CdCl_2_ was added to half the cultures and incubation continued for another 2 h. *S. aureus* strains carrying pCN47 reporter constructs were grown to early exponential (OD_540_ ∼0.15), mid-exponential (OD_540_ ∼0.8) or stationary phase (OD_540_ ∼2 after overnight culture). A sample amount corresponding to and OD_540_ of 0.5 in 1 ml was pelleted and stored at -20°C. The sample pellets were re-suspended in 100 μl digestion/lysis buffer (50 mM Tris-HCl, 20 mM MgCl_2_, 30% (w/v) raffinose) plus 1 μl of lysostaphin (12.5 μg ml^-1^) and incubated at 37°C for 1 h. Samples for SDS-PAGE were prepared by adding 4X NuPAGE LDS Sample Buffer (Invitrogen) and proteins denatured at 95°C for 10 min. Samples were separated by SDS-PAGE (NuPAGE™ 4-12% Bis-Tris Protein Gels, Invitrogen and then transferred to a PVDF transfer membrane (Thermo Scientific, 0.2 μM) following standard procedures [64, 65]. FLAG-tagged proteins were detected using mouse anti-FLAG-HRP antibody (Monoclonal ANTI-FLAG® M2-Peroxidase (HRP), Sigma-Aldrich) following the manufacturer’s protocol.

### Enzyme assay for the quantification of β-lactamase activity in transcriptional fusion plasmids

Samples for pCN41 based reporters were grown to either early exponential phase (OD-_540_∼0.15) or stationary phase for promoters with low activity (*met*C) (OD_540_ ∼2 after overnight culture) and 1 ml of culture snap frozen. β-Lactamase assays, using nitrocefin (BD Diagnostic systems) as substrate, were performed as previously described [66] using an ELx808 microplate reader (BioTek) measuring absorbance at 490 nm. Promoter activity was calculated using the following equation:

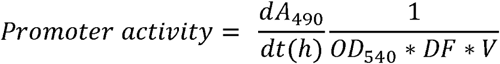

## Declarations

### Ethics approval and consent to participate

Not applicable.

### Consent for publication

Not applicable.

### Availability of data and materials

Illumina short-read sequence data was deposited at the European Nucleotide Archive with study accession number PRJEB30965. The MVF7 full genome assembly was deposited at NCBI with GenBank accession number CP062279.

### Competing interests

The authors declare that they have no competing interests.

### Funding

The study was supported by a project grant (BB/K00638X/1) and institute strategic grant funding ISP2: BB/P013740/1 from the Biotechnology and Biological Sciences Research Council (United Kingdom) to J.R.F.; grant MR/N02995X/1 from the Medical Research Council (United Kingdom) to J.R.F; Wellcome Trust collaborative award 201531/Z/16/Z to J.R.F. and J.R.P.

### Authors’ contributions

JRF and JRP conceived and led the study. GY, BAW and EJR performed the genomics analyses. AFH and MMN performed the *in vitro* analyses. SG advised on the transcriptional analyses. PH, MAT-M and R de la F provided the isolates and their metadata. GY, AFH, JRP and JRF wrote the first draft of the manuscript. All authors read and approved the final manuscript.

## Acknowledgments

We wish to thank Dr. Prerna Vohra and Dr. Cosmin Chintoan-Uta (The Roslin Institute, University of Edinburgh) for advice regarding culture of the *S. aureus* subsp. *anaerobius* isolates under microaerophilic conditions.

## Supplementary Information

**Additional file 1** (PDF format): Additional Information; Supplementary Figures S1-S6.

**Additional file 2** (XLSX format): Supplementary Tables S1-S7.

## Notes

### Competing Interest Statement

The authors have declared no competing interest.

