## Additional File 1 for "Massive genome decay and insertion sequence expansion drive the evolution of a novel host-restricted bacterial pathogen"

*[1]The Roslin Institute, University of Edinburgh, Edinburgh, United Kingdom; [2]Institute of Infection, Immunity & Inflammation, University of Glasgow, Glasgow, United Kingdom; [3]Faculty of Veterinary Medicine, University of Kufa, Kufa, Iraq; [4]Middle Euphrates Centre for Cancer and Genetic Research, University of Kufa, Kufa, Iraq; [5]Facultad de Veterinaria, Universidad Complutense de Madrid, Madrid, Spain; [6]Centre for Synthetic and Systems Biology, University of Edinburgh, Edinburgh, United Kingdom; [7]Departamento de Ciencias Biomédicas, Facultad de Ciencias de la Salud, Universidad CEU Cardenal Herrera, 46113 Moncada, Valencia, Spain; [8]Severe Infection Group, Health Research Institute Hospital La Fe, Valencia, Spain; [9]MRC Centre for Molecular Bacteriology and Infection, Imperial College London, SW7 2AZ, UK.*

[a] These authors contributed equally.

* Corresponding authors:

JRF

JRP

### Contents

**Additional information (page 2)**

**Supplementary Figures (page 7)**

**Figure S1**. Root-to-tip regression analysis**.**

**Figure S2**. Pairwise genome alignment of *S. aureus* subsp*. anaerobius versus S. aureus* subsp*. aureus*.

**Figure S3**. Phylogenetic network of representative examples of *S. aureus* Pathogenicity Islands.

**Figure S4**. Schematic representation of IS loci selected for analysis

**Figure S5.** Presence of the IS does not affect expression of the downstream gene product through active transcription.

**Figure S6**. Phylogenetic tree of the integrase gene from representative examples of *Staphylococcus aureus* prophages.

### Additional Information

#### *S. aureus* subsp. *anaerobius* MLST variability.

In contrast to previous results that ascribed all *S. aureus* subsp. *anaerobius* isolates to a single, homogenous sequence type (ST1464) [1, 2], we identified allelic variation in two of the seven housekeeping genes (*pta* and *tpi*). The four *S. aureus* subsp. *anaerobius* isolates from Sudan had an identical allelic profile, that was distinct from the one deposited in the MLST database for ST1464. Specifically, these four sequences (including the genome made available by Elbir and collaborators [3]) carried the allele 502 in the *yqiL* gene instead of allele 160 characteristic of ST1464 in the MLST database. A sequence comparison showed that these two alleles differ by one SNP. We found two *tpi* alleles in our sample set that were different in the Sudanese and European clades (alleles 177 and 422 respectively) whereas three *pta* alleles were found among the European clade. Particularly, a cluster of two Italian and one Danish samples was ascribed to ST3756, previously represented in the MLST database by a Czech sheep isolate.

#### Pseudogenes identified in *S. aureus* subsp. *anaerobius.*

In the following paragraphs, we describe some of the most noteworthy pseudogenes found according to their function, whose presence could explain the unique phenotype of *S. aureus* subsp. *anaerobius* with respect to *S. aureus* subsp. *aureus*.

***Defence mechanisms.*** We found several pseudogenised oxidoreductases (**Table S1** in Additional File 2**),** which is consistent with a deficiency in resistance to oxidative damage that reflects the microaerophilic nature of *S. aureus* subsp. *anaerobius*. The catalase gene was pseudogenised in all isolates, a feature of the *S. aureus* subsp. *anaerobius* genome already reported [4] and one of the main differences between the two subspecies of *S. aureus*. Catalase is regarded as a defensive mechanism against the oxygen radicals produced during the oxidative burst after phagocytosis, and previously, the restoration of catalase activity increased resistance to $H_2O_2$ but decreased virulence in sheep [5]. This suggests that the loss of catalase activity plays an important role in host-adaptation. Of note, genes encoding other proteins involved in resistance to oxidative killing such as *sodA* and *sodM* were intact.

Other pseudogenes related to defence mechanisms had ancestral functions associated with transmembrane pumps (ABC superfamily of transporters) and type I restriction enzymes.

***Virulence factors.*** An array of pseudogenes were associated with pathogenicity. Several were associated with the biosynthesis of the capsular polysaccharide capsule suggesting that *S. aureus* subsp. *anaerobius* may not produce a functional bacterial capsule. Others were associated with surface proteins such as (*clfB*, *spA*), and secreted proteins such as pore-forming toxins (leukocidins *lukD* and *lukE* and γ- and α-haemolysins), and the IgG-binding protein *sbi*. The coagulase gene was intact in all European isolates but present as a pseudogene in the four Sudanese isolates. Contradictory results in the literature have reported coagulase activity in isolates from Spain and Sudan but not from Kenya or France [3, 6]. All *S. aureus* subsp. *anaerobius* isolates analysed here had an intact *vwb* gene in the SaaPIMVF7 which encoded a function vWBP protein with coagulase activity.

***Metabolism and energy production.*** Twenty-nine of the 164 pseudogenes present in all isolates encoded enzymes involved in 12 amino acid metabolic pathways. Using the IMG annotation tool, we predicted that the isolate RF122 is auxotrophic for the amino acids lysine, phenylalanine, tyrosine, histidine and serine, and *Staphylococcus aureus* subsp. *anaerobius* is additionally auxotrophic, for tryptophan, arginine and leucine. Of note, the aerobic respiration pathways of *S aureus* subsp. *anaerobius* were predicted to be intact, consistent with microaerophility due to reduced resilience to oxidative stress rather than an inability to carry out aerobic respiration.

Fourteen and 6 pseudogenes encoded proteins involved in carbohydrate and lipid transport and/or metabolism, respectively. Some of the pathways affected by the presence of pseudogenes were involved in sugar metabolism (galactose, fructose, sucrose and mannose), glycolysis/gluconeogenesis, pyruvate metabolism, and phosphotransferase system (PTS, a major mechanism used for uptake of carbohydrates). One of the pseudogenes encoded the acetyl-coenzyme A synthetase, which participates in multiple distinct metabolic pathways.

Finally, production of enzymes involved in inorganic ion/coenzyme transport (specifically nickel, magnesium, manganese, cobalt, iron and molybdenum) was predicted to be affected by loss of gene function.

***S. aureus* subsp. *anaerobius* novel mobile genetic elements.**

***Prophage.*** One novel prophage (ΦSaa1) belonging to the *Siphoviridae* family was found in all isolates with a length 43.2 kb and a GC content of 33.4%, which encodes 74 proteins in most cases (some isolates presented 73 due to the absence of a HNH endonuclease). The closest prophage found by PHASTER based on nucleotide similarity (90.2%) was Φ2958PVL (accession number NC_011344), a phage initially described in methicillin-resistant *S. aureus* subsp*. aureus* isolates in Japan [7] and encoding the Panton Valentine leukocidin, which is otherwise absent in ΦSaa1 (**Fig. 1B**).

We inferred gene-by-gene homology between ΦSaa1 and Φ2958PVL using Roary [8]. Based on this comparison, 24 of the 74 genes in ΦSaa1 (32.4%) were split versions of 10 homologous genes in Φ2958PVL, whereas 37 (50%) were intact with homologues in Φ2958PVL, and 13 ΦSaa1 genes (17.6%) were exclusive. Most of the latter genes belonged to the lysogeny functional module. A phylogenetic tree constructed using FastTree v2.1.10 of integrase sequences from representative *S. aureus* subsp*. aureus* phage lineages [9] revealed that the ΦSaa1 integrase falls outside the major integrase groups (**Fig. S7** in Additional File 1), with Sa2int (the group to which the integrase of Φ2958PVL belongs) being its closest relative. The nucleotide identity between the two integrases was 78%.

***Pathogenicity island.*** We also identified one novel, 13 kb pathogenicity Island present in all isolates, which encoded 21 proteins including copies of the Staphylococcal complement inhibitor (SCIN) and the von Willebrand factor-binding protein (vWbp) (**Fig. 1B**). This *S. aureus* subsp. *anaerobius* pathogenicity island (SaaPIMVF7) is inserted in the *groES*-*groEL* site –*attB* type V, same genomic location as others such as SaPIov2 (ED133, ST133), and SaPIbov3 (RF122, ST151). We created a phylogenetic network using SplitsTree v4.14.6 and the NeighborNet algorithm [10] of SaaPIMVF7 and previously described SaPIs (**Fig. S3** in

Additional File 1). It indicated that SaaPIMVF7 is most closely related to SaPIov2 (93.1% nucleotide identity), first described in *S. aureus* subsp. *aureus* isolates of the clonal complex CC133 (responsible for infections of small ruminants including sheep and goats) and that also contains genes for vWbp and SCIN (**Fig. 1B**). A comparison of the gene homology between SaaPIMVF7 and SaPIov2 revealed that 6 of the 21 SaaPIMV7 genes (28.6%) were split versions of 3 genes found in SaPIov2, 11 (52.4%) were intact and 4 (19%) were exclusive to SaaPIMVF7. Pseudogenised genes included the genes encoding integrase and primase, suggesting SaaPIMVF7 is no longer mobile but stable in the chromosome, whereas the virulence factors vWbp and SCIN were intact. The SaaPIMVF7-encoded vWbp and SCIN proteins have 95.9% and 88.6% nucleotide identity, respectively, with the proteins encoded by SaPIov2 in *S. aureus* subsp. *aureus* strain ED133. Of note, *S. aureus* subsp. *anaerobius* also has chromosomal copies of *vwb* and *scn* of which the former is pseudogenised.

### References to Additional Information

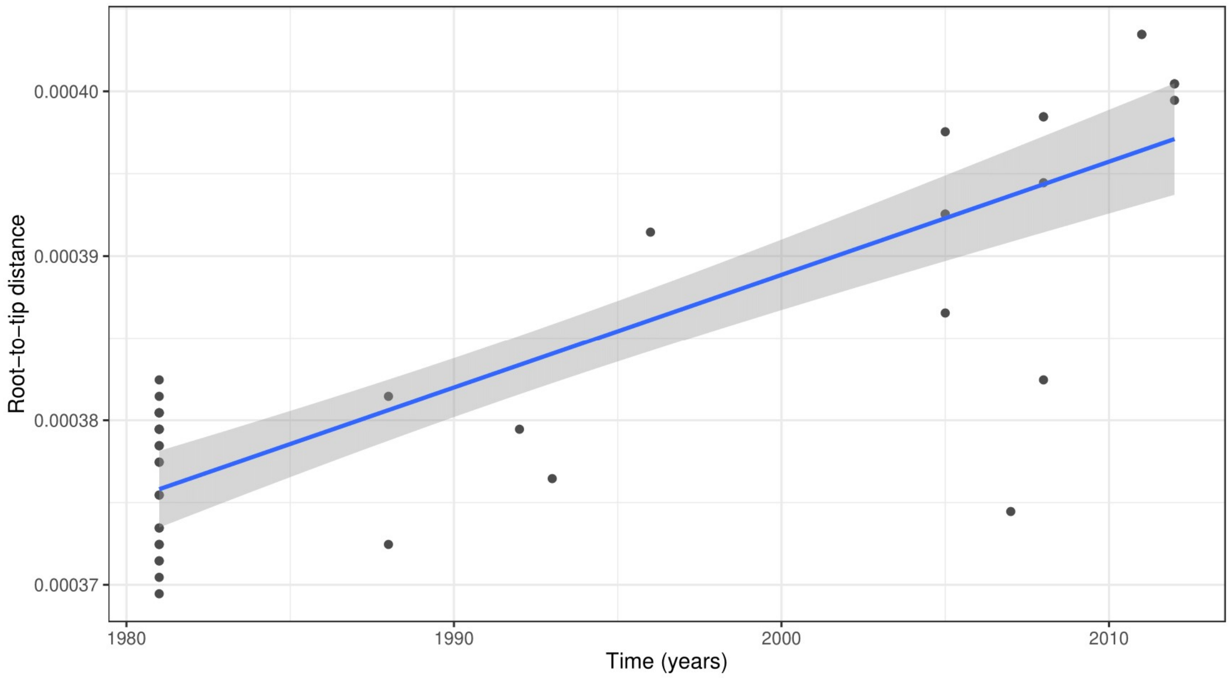


**Figure S1**. **Root-to-tip regression analysis.** Root-to-tip genetic distance against sampling time estimated from a maximum-likelihood phylogenetic tree built from a core genome alignment of *S. aureus* subsp. *anaerobius* sequences.

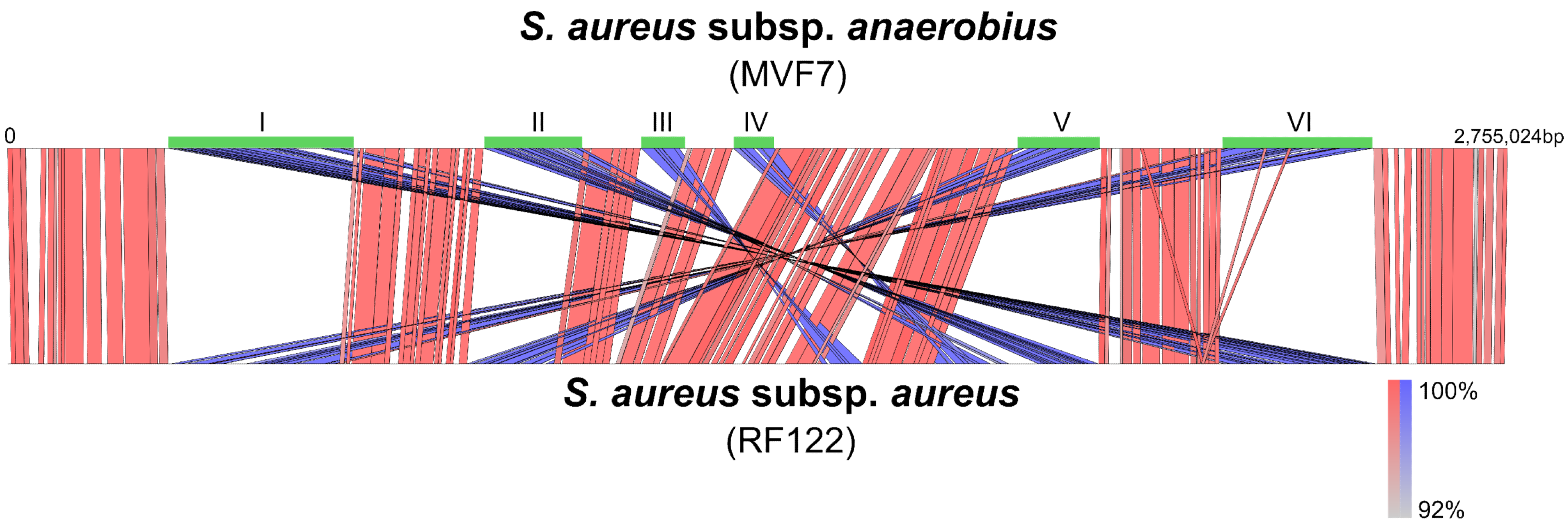


**Figure S2**. **Pairwise genome alignment of *S. aureus* subsp. *anaerobius versus S. aureus* subsp. *aureus*.** Artemis Comparison Tool (ACT) was used to compare both genomes (MVF7 and RF122, respectively). Red and blue bars indicate regions of similarity in the same and inverted orientation, respectively. The main 6 inverted chromosomal regions are highlighted in green and numbered.

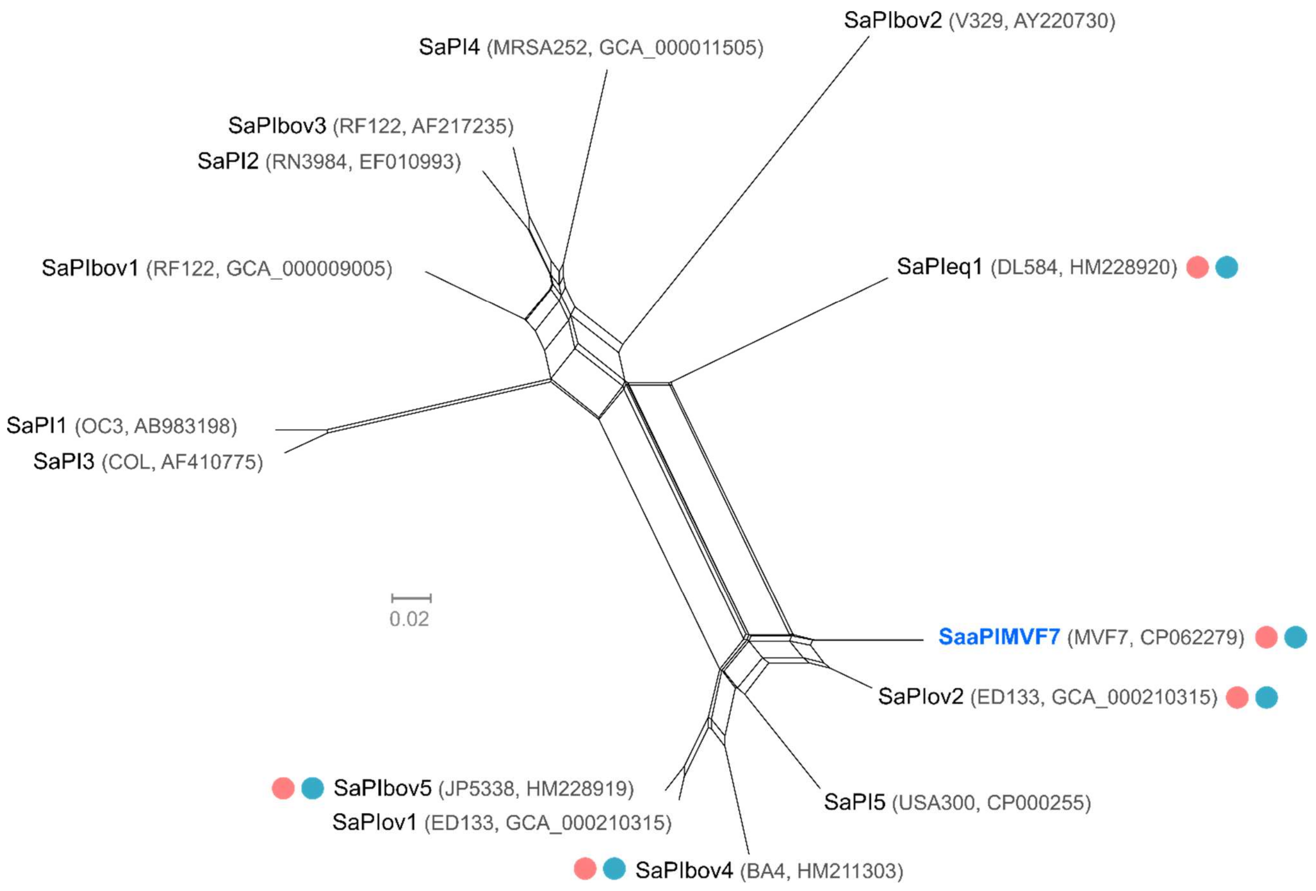


**Figure S3**. **Phylogenetic network of representative examples of *Staphylococcus aureus* Pathogenicity Islands.** In bold and blue the SaPI found in *S. aureus* subsp. *anaerobius* (SaaPIMVF7). The red and blue circles indicate those SaPIs that harbour the genes *vwb* and *scn*. Reference sequences are labelled indicating SaPI name, isolate and accession number (of the SaPI sequence when available, of the original genome otherwise).

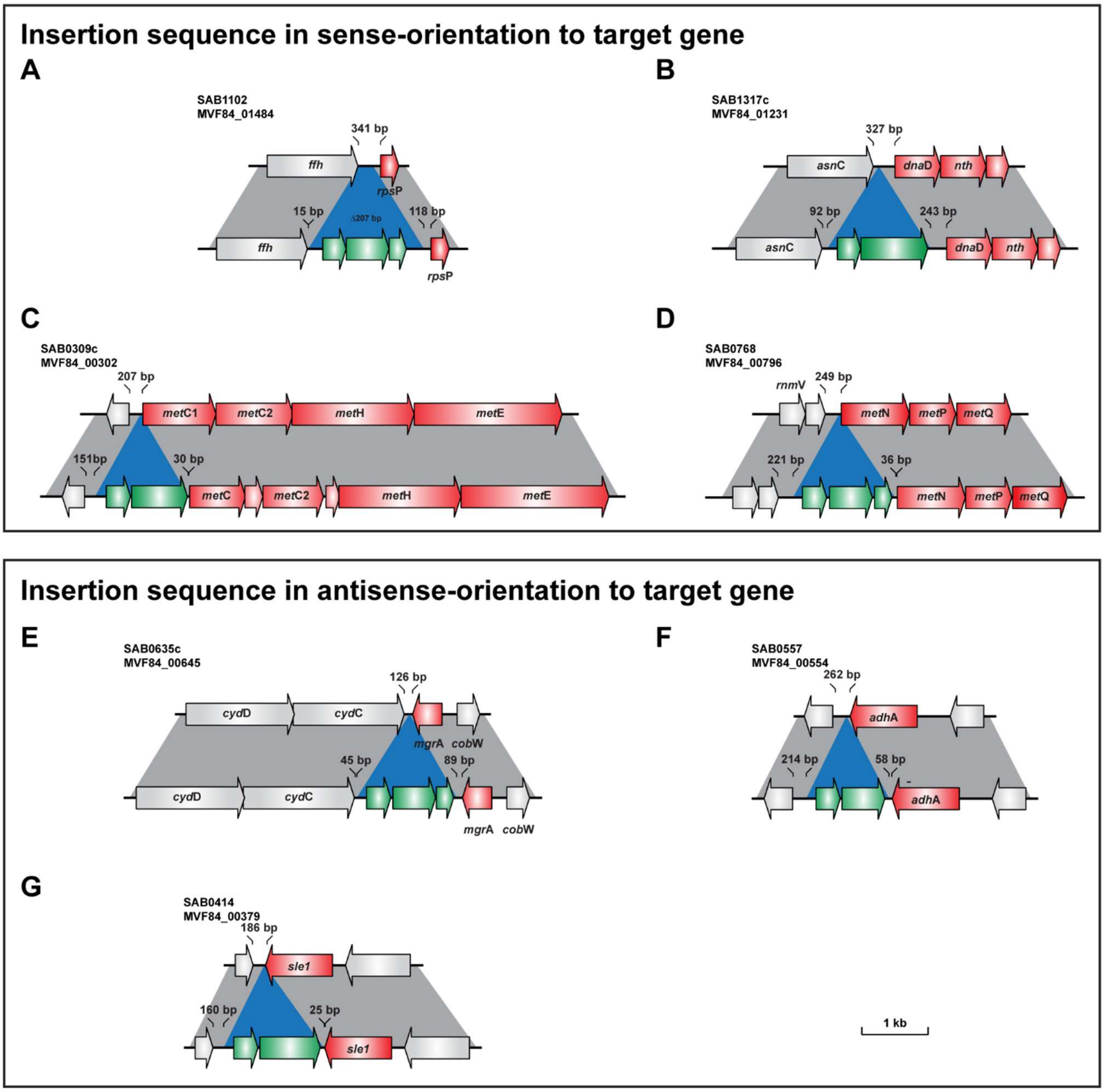


**Figure S4. Schematic representation of IS loci selected for analysis.** IS loci are shown for *S. aureus* subsp._*anaerobius* strain MVF84 and *S. aureus* strain RF122 representing the ancestral genomic context. (A-D) IS inserted at various distances from the downstream gene start codon. Note that in (A) IS insertion results in a 207 bp deletion in the intergenic region in strain MVF84 relative to strain RF122. (E-G) IS inserted downstream and in antisense orientation of target gene. (E&G) Locus in RF122 shows antisense orientation of downstream gene while in (F) downstream gene is in the same orientation as target gene for IS.

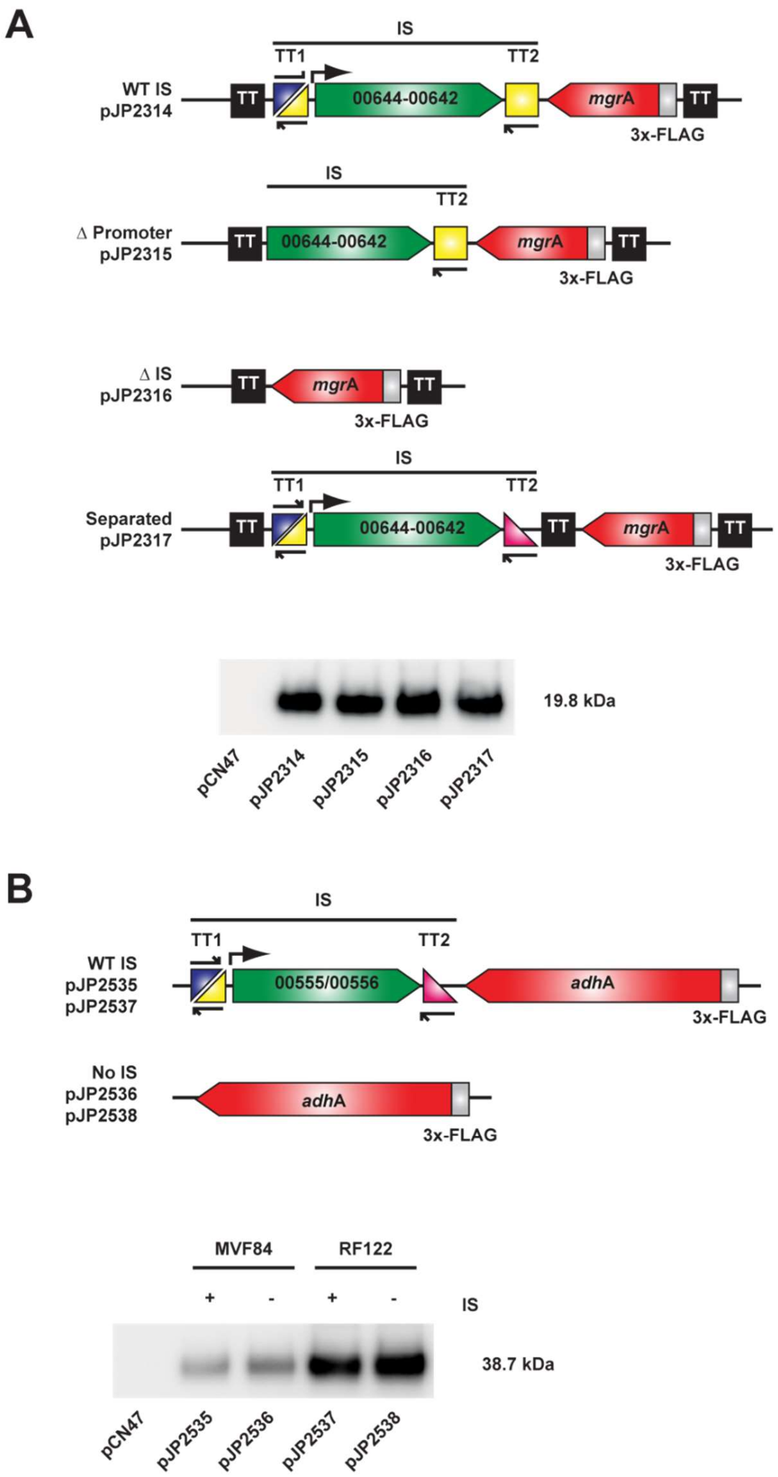


**Figure S5. Presence of the insertion sequence (IS) does not affect expression of the downstream gene product through active transcription.** Western blot analysis of the depicted expression constructs for assessing the impact of IS on the expression of (A) MgrA or (B) AdhA from the IS encoded promoter. 3x-FLAG-tagged protein-encoding genes containing or missing the IS were cloned into pCN47 and plasmids introduced into the *S. aureus* subsp. *aureus* strain RN4220 Δ*spa* for analysis. For a schematic of the locus in either *S. aureus* subsp. *anaerobius* MVF84 or *S. aureus* subsp. *aureus* RF122 refer to Figure S4.

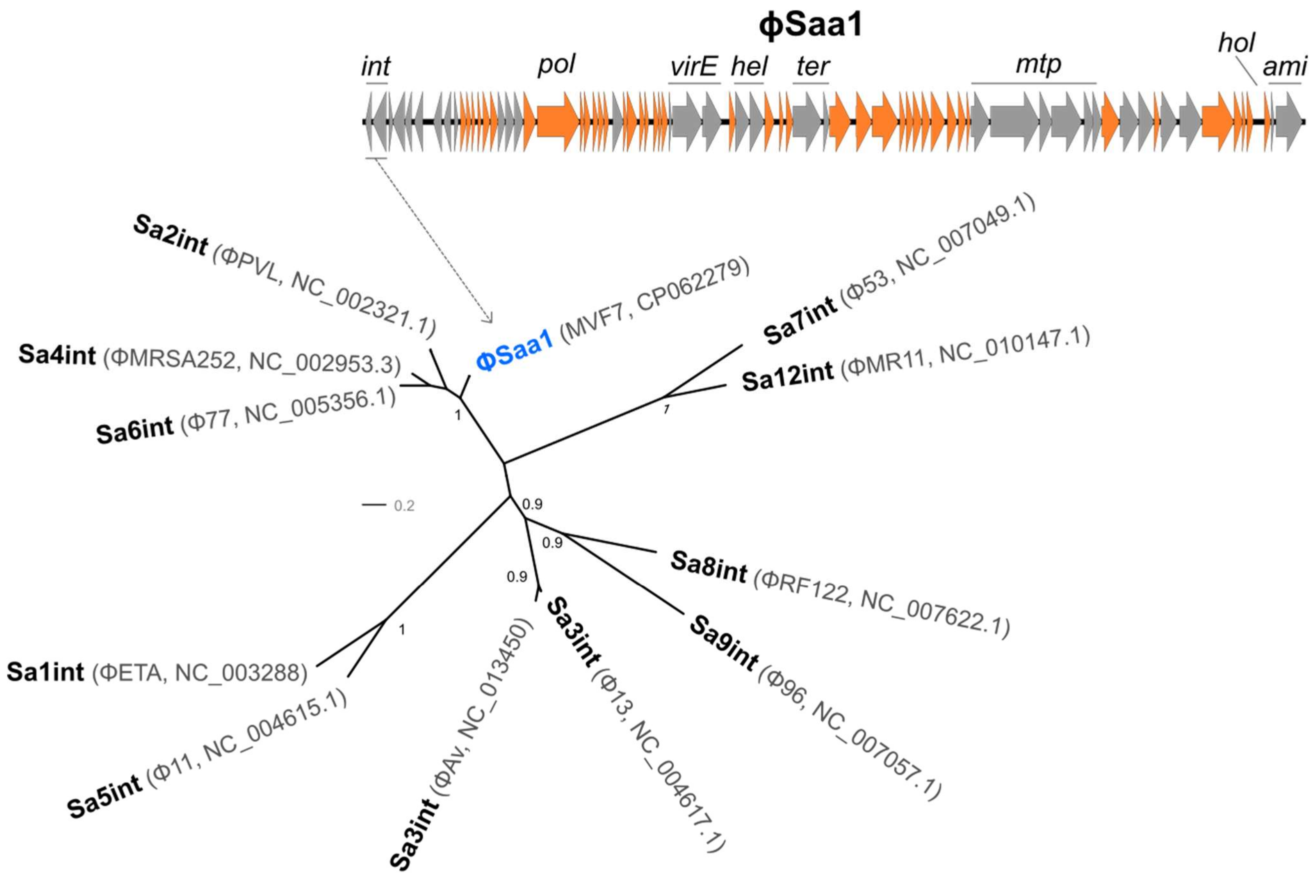


**Figure S6**. **Phylogenetic tree of the integrase gene from representative examples of *S. aureus* prophages**. Top: genome map of the phage found in *S. aureus* subsp. *anaerobius* (ΦSaa1). Genes in grey are pseudogenes and genes in orange are intact by comparison to Φ2958PVL. *Int:* integrase; *pol:* polymerase; *virE:* virulence protein E; *hel:* helicase; *ter:* terminase; *mtp:* measure tape protein; *hol:* holin; *ami:* amidase. Bottom: integrase tree. In bold and blue the integrase of ΦSaa1. Reference sequences are labelled indicating integrase major group, phage and accession number.
